# Probabilistic Count Matrix Factorization for Single Cell Expression Data Analysis

**DOI:** 10.1101/211938

**Authors:** G. Durif, L. Modolo, J. E. Mold, S. Lambert-Lacroix, F. Picard

## Abstract

**Motivation:** The development of high throughput single-cell sequencing technologies now allows the investigation of the population diversity of cellular transcriptomes. The expression dynamics (gene-to-gene variability) can be quantified more accurately, thanks to the measurement of lowly-expressed genes. In addition, the cell-to-cell variability is high, with a low proportion of cells expressing the same genes at the same time/level. Those emerging patterns appear to be very challenging from the statistical point of view, especially to represent a summarized view of single-cell expression data. PCA is a most powerful tool for high dimensional data representation, by searching for latent directions catching the most variability in the data. Unfortunately, classical PCA is based on Euclidean distance and projections that poorly work in presence of over-dispersed count data with dropout events like single-cell expression data.

**Results:** We propose a probabilistic Count Matrix Factorization (pCMF) approach for single-cell expression data analysis, that relies on a sparse Gamma-Poisson factor model. This hierarchical model is inferred using a variational EM algorithm. It is able to jointly build a low dimensional representation of cells and genes. We show how this probabilistic framework induces a geometry that is suitable for single-cell data visualization, and produces a compression of the data that is very powerful for clustering purposes. Our method is competed against other standard representation methods like t-SNE, and we illustrate its performance for the representation of single-cell expression (scRNA-seq) data.

**Availability:** Our work is implemented in the pCMF R-package^1^.

## 1 Introduction

The combination of massive parallel sequencing with high-throughput cell biology technologies has given rise to the field of single-cell genomics, which refers to techniques that now provide genome-wide measurements of a cell’s molecular profile either based on DNA (Zong *et al*., 2012), RNA (Picelli *et al*., 2013), or chromatin (Buenrostro *et al*., 2015; Rotem *et al*., 2015). Similar to the paradigm shift of the 90s characterized by the first molecular profiles of tissues (Golub *et al*., 1999), it is now possible to characterize molecular heterogeneities at the cellular level (Deng *et al*., 2014; Saliba *et al*., 2014). A tissue is now viewed as a population of cells of different types, and many fields have now identified intra-tissue heterogeneities, in T cells (Buettner *et al*., 2015), lung cells (Trapnell *et al*., 2014), or intestine cells (Grün *et al*., 2015). The construction of a comprehensive atlas of human cell types is now within our reach (Wagner *et al*., 2016). The characterization of heterogeneities in single-cell expression data thus requires an appropriate statistical model, as the transcripts abundance is quantified for each cell using read counts. Hence, standard methods based on Gaussian assumptions are likely to fail to catch the biological variability of lowly expressed genes, and Poisson or Negative Binomial distributions constitute an appropriate framework (Chen *et al*., 2016; Riggs and Lalonde, 2017, Chap. 6). Moreover, dropouts, either technical (due to sampling difficulties) or biological (no expression or stochastic transcriptional activity), constitute another major source of variability in scRNA-seq (single-cell RNA-seq) data, which has motivated the development of the so-called Zero-Inflated models (Kharchenko *et al*., 2014).

Principal component analysis (PCA) is one of the most widely used dimension reduction technique, as it allows the quantification and visualization of variability in massive datasets. It consists in approximating the observation matrix **X**_[*n×m*]_ (*n* cells, *m* genes), by a factorized matrix of reduced rank, denoted **UV**^*T*^ where **U**_[*n×K*]_ and **V**_[*m×K*]_ represent the latent structure in the observation and variable spaces respectively. This projection onto a lower-dimensional space (of dimension *K*) allows one to catch gene co-expression patterns and clusters of individuals. PCA can be viewed either geometrically or through the light of a statistical model (Landgraf and Lee, 2015). Standard PCA is based on the *ℓ*_2_ distance as a metric and is implicitly based on a Gaussian distribution (Eckart and Young, 1936). Model-based PCA offers the unique advantage to be adapted to the data distribution. It consists in specifying the distribution of the data **X**_[*n×m*]_ through a statistical model, and to factorize 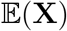 instead of **X**. In this context the *ℓ*_2_ metric is replaced by the Bregman divergence which is adapted to maximum likelihood inference (Collins *et al*., 2001). A probabilistic version of the Gaussian PCA was proposed by Pierson and Yau (2015) in the context of single cell data analysis, with the modeling of zero inflation (the ZIFA method). ScRNA-seq data may be better analyzed by methods dedicated to count data such as the Non-negative Matrix Factorization (NMF) introduced in a Poisson-based framework by Lee and Seung (1999) or the Gamma-Poisson factor model (Cemgil, 2009; Févotte and Cemgil, 2009; Landgraf and Lee, 2015). None of the currently available dimension reduction methods fully model single-cell expression data, characterized by over-dispersed zero inflated counts (Kharchenko *et al*., 2014; Zappia *et al*., 2017).

Our method is based on a probabilistic count matrix factorization (pCMF). We propose a dimension reduction method that is dedicated to over-dispersed counts with dropouts, in high dimension. In particular, gene expression can be normalized but does not require to be transformed (log, Anscombe) in our framework. Our factor model takes advantage of the Poisson Gamma representation to model counts from scRNA-seq data (Zappia *et al*., 2017). In particular, we use Gamma priors on the distribution of principal components. We model dropouts with a Zero-Inflated Poisson distribution (Simchowitz, 2013), and we introduce sparsity in the model thanks to a spike-and-slab approach (Malsiner-Walli and Wagner, 2011) that is based on a two component sparsity-inducing prior on loadings (Titsias and Lázaro-Gredilla, 2011). We propose a heuristic to initialize the sparsity layer based on the variance of the recorded variables, acting as an integrated gene filtering step, which is an important issue in scRNA-seq data analysis (Soneson and Robinson, 2018). The model is inferred using a variational EM algorithm that scales favorably to data dimension compared with Markov Chain Monte-Carlo (MCMC) methods (Hoffman *et al*., 2013; Blei *et al*., 2017). Then we propose a new criterion to assess the quality of fit of the model to the data, as a percentage of explained deviance, following a strategy that is standard in the Generalized Linear Models framework. Moreover, we show that our criterion corresponds to the percentage of explained variance in the PCA case, which makes it suitable to compare geometric and probabilistic methods.

We show the performance of pCMF on simulated and experimental datasets, in terms of visualization and quality of fit. Moreover, we show the benefits of using pCMF as a preliminary dimension reduction step before clustering or before the popular t-SNE approach (van der Maaten and Hinton, 2008; Amir *et al*., 2013). Experimental published data are used to show the capacity of pCMF to provide a better representation of the heterogeneities within scRNA-Seq datasets, which appears to be extremely helpful to characterize cell types. Finally, our approach also provides a lower space representation for genes (and not only for cells), contrary to t-SNE. pCMF is implemented in the form of a R package available at https://github.com/gdurif/pCMF.

## 2 Count Matrix Factorization for zero-inflated overdispersed data

### The Poisson factor model

Our data consist of a matrix of counts (potentially normalized but not transformed), denoted by 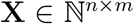, that we want to decompose onto a subspace of dimension *K* (being fixed). In a first step we suppose that the data follow a multivariate Poisson distribution of intensity **Λ**. Following the standard Poisson Non-Negative Matrix Factorization (Poisson NMF, Lee and Seung, 1999), we approximate this intensity such that

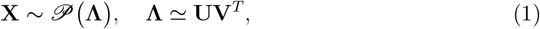

with factor **U** ∈ ℝ^+,*n×K*^ the coordinates of the *n* observations (cells) in the subspace of dimension *K*, and loadings **V** ∈ ℝ^+,*m×K*^ the contributions of the *m* variables (genes).

### Modeling over-dispersion

We account for over-dispersion by using the the Negative-Binomial distribution (Anders and Huber, 2010), through a hierarchical Gamma-Poisson representation (GaP) Cemgil (2009). In our factor model **U** and **V** are modeled as independent random latent variables with Gamma distributions such that

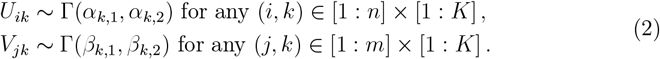

In practice, some third-party latent variables are introduced for the derivation of our inference algorithm (Cemgil, 2009; Zhou *et al*., 2012). We consider latent variables **Z** = [*Z_ijk_*] ∈ ℝ^*n×m×K*^, defined such that *X_ij_* = Σ*_k_ Z_ijk_*. These new indicator variables quantify the contribution of factor *k* to the data. Here *Z_ijk_* are assumed to be conditionally independent and to follow a conditional Poisson distribution, i.e. 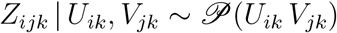. Thus, the conditional distribution of *X_ij_* remains 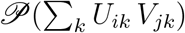 thanks to the additive property of the Poisson distribution.

### Dropout modeling with a zero-inflated (ZI) model

To model zero-inflation, i.e. random null observations called dropout events, we introduce a dropout indicator variable *D_ij_* ∈ {0,1} for *i* = 1,…,*n* and *j* = 1,…,*p* (c.f. Simchowitz, 2013). In this context, each *D_ij_* = 0 if gene *j* has been subject to a dropout event in cell *i*, with 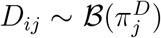. We consider gene-specific dropout rates, 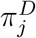, following recommendations of the literature (Pierson and Yau, 2015). Thus, to include zero-inflation in the probabilistic factor model, we consider that

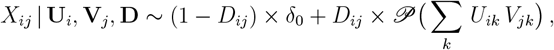

where *δ*_0_ is the Dirac mass at 0, i.e. *δ*_0_(*X_ij_*) = 1 if *X_ij_* = 0 and 0 otherwise. The dropout indicators *D_ij_* are assumed to be independent from the factors *U_ik_* and *V_jk_*. Then, by integrating *D_ij_* out, the probability of observing a zero in the data illustrates the two potential sources of zeros and becomes

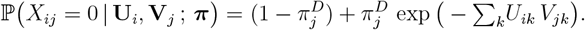

Thus, inference will be based on the factors *U_ik_* and *V_jk_* and on probabilities 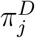.

### Probabilistic variable selection

Finally we suppose that our model is parsimonious. We consider that among all recorded variables, only a proportion carries the signal and the others are noise. To do so, we modify the prior of the loadings variables *V_jk_*, to consider a sparse model with a two-group sparsity-inducing prior (Engelhardt and Adams, 2014). The model is then enriched by the introduction of a new indicator variable 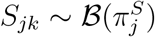, that equals 1 if gene *j* contributes to loading *V_jk_*, and zero otherwise. 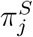 stands for the prior probability for gene *j* to contribute to any loading. To define the sparse GaP factor model, we modify the distribution of the loadings latent factor *V_jk_*, such that

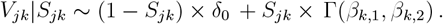

This spike-and-slab formulation (Mitchell and Beauchamp, 1988) ensures that *V_jk_* is either null (gene *j* does not contribute to factor *k*), or drawn from the Gamma distribution (when gene *j* contributes to the factor). The contribution of gene *j* to the component *k* is accounted for in the conditional Poisson distribution of *X_ij_*, with

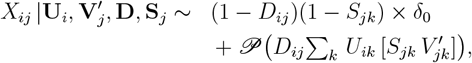

where 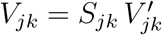 such that 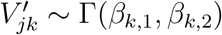.

### Underlying geometry

Knowing **U** and **V**, to quantify the approximation of matrix **X** by **UV**^*T*^, we consider the Bregman divergence, that can be viewed as a generalization of the Euclidean metric to the exponential family (see Collins *et al*., 2001; Banerjee *et al*., 2005; Chen *et al*., 2008). In the Poisson model, the Bregman divergence between **X** and **UV**^*T*^ is defined as (Févotte and Cemgil, 2009):

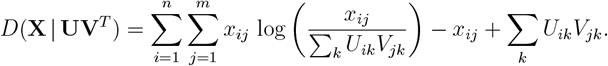

Hence the geometry is induced by an appropriate probabilistic model dedicated to count data. Potential identifiability issues are addressed in Supp. Mat. (Section S.2).

In the following, we will refer to pCMF for the method implementing the model with dropout but the without sparsity layer, and to sparse pCMF (or spCMF) for the model with dropout and sparsity layers.

### 2.1 Quality of the reconstruction

The Bregman divergence between the data matrix **X** and the reconstructed matrix 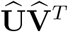 in our GaP factor model is related to the deviance of the Poisson model defined such as (Landgraf and Lee, 2015)

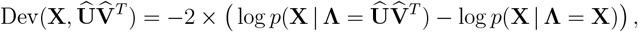

where log *p*(**X|Λ**) is the Poisson log-likelihood based on the matrix notation (1). We have 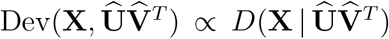, thus the deviance can be used to quantify the quality of the model.

Regarding PCA, the percentage of explained variance is a natural and unequivocal quantification of the quality of the representation. We introduce a criterion that we call percentage of explained deviance that is a generalization of the percentage of explained variance to our GaP factor model. However, since our models are not nested for increasing *K*, the definition of this criterion appears non trivial. To assess the quality of our model, we propose to define the percentage of explained deviance as:

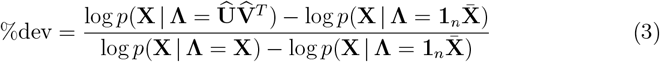

where 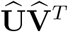 is the predicted reconstructed matrix in our model, 1_*n*_ is a column vector filled with 1 and 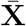 is a row vector of size *m* storing the column-wise average of **X**. We use two baselines: (*i*) the log-likelihood of the saturated model, i.e. log *p*(**X|Λ** = **X**) (as in the deviance), which corresponds to the richest model and (*ii*) the log-likelihood of the model where each Poisson intensities λ_*ij*_ is estimated by the average of the observations in the column *j*, i.e. 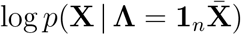, which is the most simple model that we could use. This formulation ensures that the ratio %dev lies in [0; 1]. An interesting point is that if we assume a Gaussian distribution on the data, the percentage of explained deviance is exactly the percentage of explained variance from PCA (c.f. Section S.1), which makes our criterion suitable for to compare different factor models.

### 2.2 Choosing the dimension of the latent space

As noticed in the previous section, the GaP factor model with an increasing number *K* of factors are not nested (the model associated to the NMF presents the same properties). Consequently, testing different values of *K* requires to fit different models (contrary to PCA for instance). We choose the number of factors by fitting a model with a large *K* and verifying how the matrix 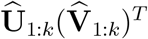 reconstructs **X** depending on *k* = 1,…,*K* with a rule of thumb based on the “elbow” shape of the fitting criterion. This approach is for instance widely used in PCA by checking the proportion of variability explained by each components, see Friguet (2010, p.96) for a review of the different criteria to choose *K* in this context. Here we use the deviance, or equivalently the Bregman divergence 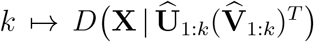 to find the latent dimension from where adding new factors does not improve 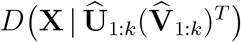. This determination is however not always unambiguous and may sometimes lead to some over-fitting, i.e. when considering too much factors. In addition, when focusing on data visualization, we generally set *K* = 2.

## 3 Model inference using a variational EM algorithm

Our goal is to infer the posterior distributions over the factors **U** and **V** depending on the data **X**. To avoid using the heavy machinery of MCMC (Nathoo *et al*., 2013) to infer the intractable posterior of the latent variables in our model, we use the framework of variational inference (Hoffman *et al*., 2013). In particular, we extend the version of the variational EM algorithm (Beal and Ghahramani, 2003) proposed by Dikmen and Févotte (2012) in the context of the standard Gamma-Poisson factor model to our sparse and zero-inflated GaP model. Figure S.1 in Supp. Mat. gives an overview of the variational framework.

### 3.1 Definition of variational distributions

In the variational framework, the posterior *p*(**Z, U, V′, S, D|X**) is approximated by the variational distribution *q*(**Z, U, V′, S, D**) regarding the Kullback-Leibler divergence (Hoffman *et al*., 2013), that quantifies the divergence between two probability distributions. Since the posterior is not explicit, the inference of *q* is based on the optimization of the Evidence Lower Bound (ELBO), denoted by *J*(*q*) and defined as:

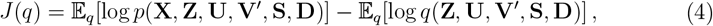

that is a lower bound on the marginal log-likelihood log *p*(**X**). In addition, maximizing the ELBO *J*(*q*) is equivalent to minimizing the KL divergence between *q* and the posterior distribution of the model (Hoffman *et al*., 2013). To derive the optimization, *q* is assumed to lie in the mean-field variational family, i.e. (*i*) to be factorisable with independence between latent variables and between observations and (*ii*) to follow the conjugacy in the exponential family, i.e. to be in the same exponential family as the full conditional distribution on each latent variables in the model. Thanks to the first assumption, in our model, the variational distribution *q* is defined as follows:

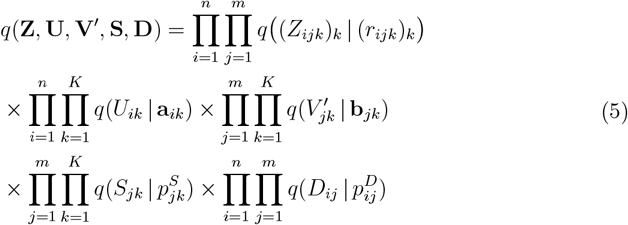

where 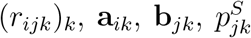 and 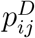 are the parameters of the variational distribution regarding 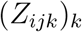, *U_ik_*, 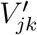, *S_jk_*, *D_ij_* respectively. Then we need to precise the full conditional distributions of the model before defining the variational distributions more precisely.

### 3.2 Approximate posteriors

To approximate the (intractable) posterior distributions, variational distributions are assumed to lie in the same exponential family as the corresponding full conditionals and to be independent such that:

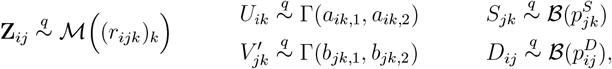

where 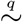 denotes the variational distribution. The strength of our approach is the resulting explicit approximate distribution on the loadings that induces sparsity:

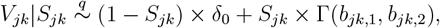

In the following, the derivation of variational parameters involves the moments and log-moments of the latent variables regarding the variational distribution. Since the distributions q is fully determined, these moments can be directly computed. For the sake of simplicity, we will use notation 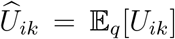 and 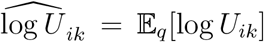 (collected in the matrices 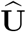 and 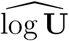 respectively), with similar notations for other hidden variables of the model 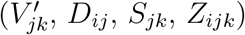.

### 3.3 Derivation of variational parameters

In order to find a stationary point of the ELBO, *J*(*q*) is differentiated regarding each variational parameter separately. The formulation of the ELBO regarding each parameter separately is based on the corresponding full conditional, e.g. *p*(*U_ik_*|—), *p*(*V_jk_*|—) and *p*((*Z_ijk_*)_*k*_|—). The partial formulation are therefore respectively:

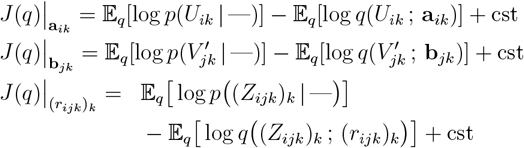

Similar formulations can be derived regarding parameters 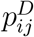 and 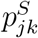. Therefore, the ELBO is explicit regarding each variational parameter and the gradient of the ELBO *J*(*q*) depending on the variational parameters **a**_*ik*_, **b**_*jk*_, *r_ijk_*, 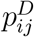 and 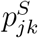 respectively can be derived to find the coordinate of the stationary point (corresponding to a local optimum). In our factor model all full conditionals are tractable (c.f. Section S.4.1 in Supp. Mat.). In practice, thanks to the formulation in the exponential family, the optimum value for each variational parameter corresponds to the expectation regarding *q* of the corresponding parameter of the full conditional distribution (see Hoffman *et al*., 2013). Thus the coordinates of the ELBO’s gradient optimal point are explicit.

We mentioned that distributions with a mass at 0 (zero-inflated Poisson or sparse Gamma) lie in the exponential family (Eggers, 2015) and the general formulation from Hoffman *et al*. (2013) remains valid. Detailed formulations of update rules regarding all variational parameters are given in Supp. Mat. (Section S.4.2).

### 3.4 Variational EM algorithm

We use the variational-EM algorithm (Beal and Ghahramani, 2003) to jointly approximate the posterior distributions and to estimate the hyper-parameters **Ω** = (***α, β, π^S^, π^D^***).

In this framework, the variational inference is used within a variational E-step, in which the standard expectation of the joint likelihood regarding the posterior 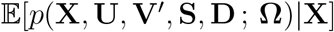 is approximated by 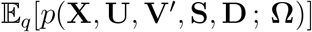. Then the variational M-step consists in maximizing 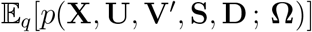 w.r.t. the hyper-parameters **Ω**. In the variational-EM algorithm, we have explicit formulations of the stationary points regarding variational parameters (E-step) and prior hyper-parameters (M-step) in the model, thus we use a coordinate descent iterative algorithm (see Wright, 2015, for a review) to infer the variational distribution. Detailed formulations of update rules regarding all prior hyper-parameters are given in Supp. Mat. (Section S.4.3).

### 3.5 Initialization of the algorithm

To initialize variational and hyper-parameters of the model, we sample **U** and **V** from Gamma distributions such that *X_ij_* ≃ Σ_*k*_ *U_ik_V_jk_* on average. The Gamma variational and hyper parameters are initialized from these values following the update rules detailed in Supp. Mat. Section S.4.2. Dropout probabilities 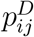 and 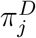 are initialized by 1/*n*Σ_*i*_ 1_{*X_ij_*>0}_, i.e. the proportion of non-zero in the data for the corresponding gene. To initialize the sparsity probabilities 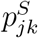 and 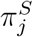, we use a heuristic based on the variance of the corresponding gene *j*. Assuming that genes with low variability will have less impact on the structure embedded in the data, we propose a starting value such that

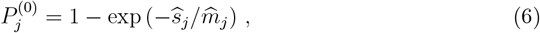

where 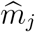 is the mean of the non-null observations for gene *j* and 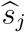 its standard deviation (including null values). The scaling is better when removing the null values to compute the mean. This quantity adapts to the empirical variance of the observations, and will be close to 0 for genes with low variance, and close to 1 for genes with high variability.

## 4 Empirical study of pCMF

All codes are available on a public repository for reproducibility^2^. We compare our method with standard approaches for unsupervised dimension reduction: the Poisson-NMF (Lee and Seung, 1999), applied to raw counts (model-based matrix factorization approach based on the Poisson distribution); the PCA (Pearson, 1901) and the sparse PCA (Witten *et al*., 2009), based on an *ℓ*_1_ penalty in the optimization problem defining the PCA to induce sparsity in the loadings **V**, both applied to log counts. We use sparse methods (sparse PCA, sparse pCMF) with a re-estimation step on the selected genes. We will refer to them as (s)PCA and (s)pCMF in the results respectively, to differentiate them from sparse PCA and sparse pCMF (without re-estimation), PCA and pCMF (without the sparsity layer). In addition, we use the Zero-Inflated Factor Analysis (ZIFA) by Pierson and Yau (2015), a dimension reduction approach that is specifically designed to handle dropout events in single-cell expression data (based on a zero-inflated Gaussian factor model applied to log-transformed counts). We present quantitative clustering results and qualitative visualization results on simulated and experimental scRNA-seq data. We also compare our method with t-SNE that is commonly used for data visualization (van der Maaten and Hinton, 2008). It requires to choose a “perplexity” hyper-parameter that cannot be automatically calibrated, thus being less appropriate for a quantitative analysis. In the following, we always choose the perplexity values that gives the better clustering results.

### 4.1 Simulated data analysis

To generate synthetic data we follow the hierarchical Gamma-Poisson framework as adopted by others (Zappia *et al*., 2017). Details are provided in the Supp. Mat. (Section S.5). We generate synthetic multivariate over-dispersed counts, with *n* = 100 individuals and *m* = 800 recorded variables. We artificially create clusters of individuals (with different level of dispersion) and groups of dependent variables. We we set different levels of zero-inflation in the data (i.e. low or high probabilities of dropout events, corresponding to random null values in the data), and some part of the m variables are generated as random noise that do not induce any latent structure. Thus, we can test the performance of our method in different realistic data configurations, the range of our simulation parameters being comparable to other published simulated data (Zappia *et al*., 2017).

We train the different methods with *K* = 2 to visualize the reconstructed matrices 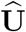 and 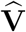 (c.f. Section 2). To assess the ability of each method to retrieve the structure of cells or genes, we run a *κ*-mean clustering algorithm on 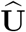 and 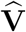 respectively (with *κ* = 2) and we measure the adjusted Rand Index (Rand, 1971) quantifying the accordance between the predicted clusters and original groups of cells or genes. Regarding our approach pCMF, we use 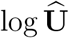 and 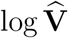 for data visualization and clustering because the log is the canonical link function for Gamma models. In addition, we also compute the percentage of explained deviance associated to the model to assess the quality of the reconstruction. Regarding the PCA (sparse or not) and ZIFA, we use the standard explained variance criterion (c.f. Section 2.1).

#### 4.1.1 Clustering in the observation space

##### Effect of zero-inflation and cell representation

We first assess the robustness of the different methods to the level of dropout or zero-inflation (ZI) in the data. We generate data with 3 groups of observations (c.f. Supp. Mat. Section S.5) with a wide range of dropout probabilities. Figure 1a shows that (s)pCMF adapts to the level of dropout in the data and recovers the original clusters (high adjusted Rand Index) even with dropouts. Despite comparable performance with low dropout, Poisson-NMF and ZIFA are very sensitive to the addition of zeros. In addition, methods based on transformed counts like (s)PCA and t-SNE perform poorly, as they do not account for the specificity of the data (discrete, over-dispersed, (O’Hara and Kotze, 2010)).

**Figure 1:**
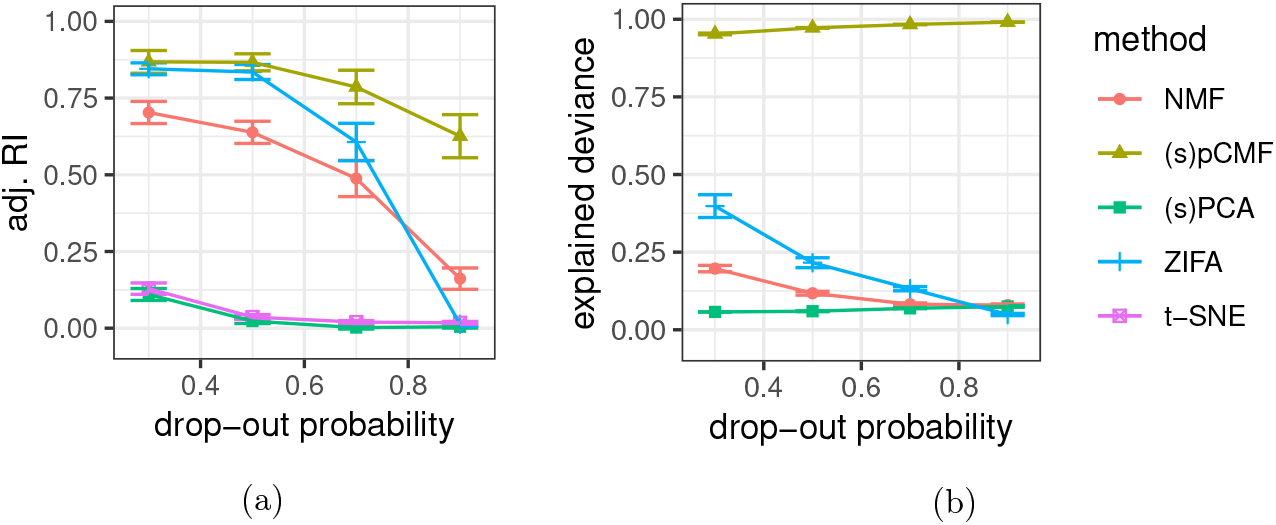
Adjusted Rand Index (1a) for the clustering on 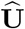 versus the true groups of cells; and explained deviance (1b) depending on the probability used to generate dropout events. Average values and deviation are estimated across 50 repetitions.

##### Effect of noisy genes and gene representation

To quantify the impact of noisy genes on the retrieval of the clusters, we consider data generated with different proportions of noisy genes (genes that do not induce any structure in the data). We generate data with three groups of genes: two groups inducing some latent structure and one group of noisy genes (c.f. Supp. Mat. Section S.5). Figure 2 shows that (s)pCMF identifies correctly the clusters of genes, including the set of noisy genes, contrary to other approaches. This point shows that our approach correctly identifies the genes that support the lower-dimensional representation.

**Figure 2:**
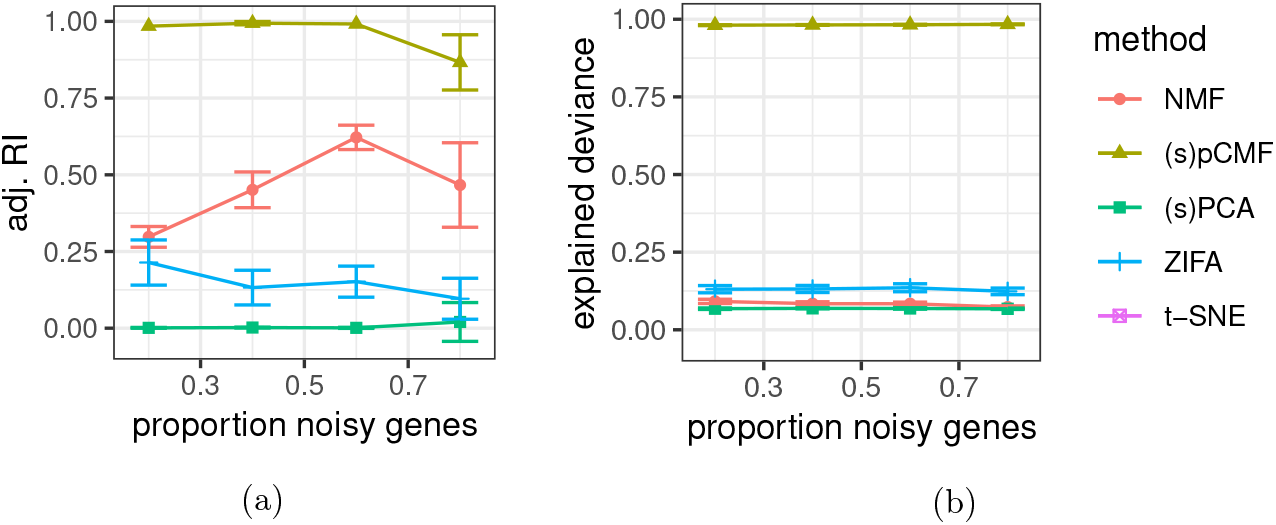
Adjusted Rand Index (2a) for the clustering on 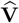 versus the true groups of genes; and explained deviance (2b) depending on the proportion of noisy genes. Average values and deviation are estimated across 50 repetitions.

In addition to the clustering results, we compared the selection accuracy of the only two methods (sPCA, spCMF) that perform variable selection (Supp. Mat. Figure S.2). A selected gene is a gene that contributes to any latent dimension (any *V_jk_* ≠ 0). Sparse pCMF performs better than sparse PCA for various latent dimension *K* even for high levels of noisy genes. Sparse PCA shows better selection accuracy when the proportion of noisy genes is low. This point suggests that sparse pCMF would be less sensitive to gene pre-filtering when analysing scRNA-seq data, which corresponds to a removal of noisy genes and is generally crucial (Soneson and Robinson, 2018).

Details about data generation and additional data configuration regarding Figures 1 and 2 can be found in Supp. Mat. (Section S.7, especially Figures S.3 and S.4).

#### 4.1.2 Data visualization

Data visualization is central in single-cell transcriptomics for the representation of high dimensional data in a lower dimensional space, in order to identify groups of cells, or to illustrate the cells diversity (e.g. Llorens-Bobadilla *et al*., 2015; Segerstolpe *et al*., 2016). In the matrix factorization framework, we represent observation (cell) coordinates and variable (gene) contributions: resp. 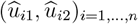 and 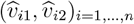 (or their log transform for pCMF) when the dimension is *K* = 2 (see Section 2).

We consider the same simulated data as previously (*n* =100, *m* = 800, with three groups of cells, two groups of relevant genes and the set of noisy genes). Our visual results are consistent with the previous clustering results both regarding cell and gene visualization. In this challenging context (high zero-inflation and numerous noisy variables), by using pCMF, we are able to graphically identify the groups of individuals (cells) in the simulated zero-inflated count data (c.f. Figure 3). On the contrary, the 2-D visualization is not successful with PCA, ZIFA, Poisson-NMF and t-SNE, illustrating the interest of our data-specific approach. This point supports our claim that using data-specific model improves the quality of the reconstruction in the latent space.

**Figure 3:**
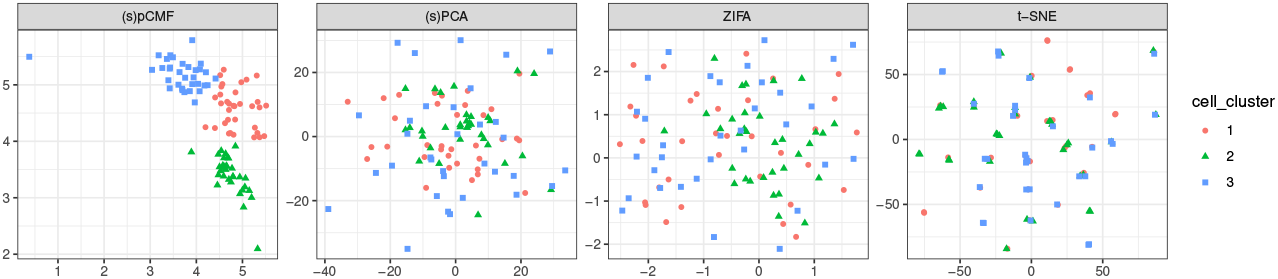
Representation of cells in a subspace of dimension *K* = 2. Here we have 60% of noisy variables, and dropout probabilities around 0.9.

In addition, linear projection methods (NMF, PCA, pCMF, ZIFA) can be used to visualize the contribution of genes to the principal axes (c.f. Figure 4). Thanks to sparsity constraints, the contribution of noisy genes are mostly set to 0 for sparse pCMF. Surprisingly, this selection is not efficient in the case of sparse PCA, indicating a lack of calibration of the sparsity constraint when data are counts. In comparison, Poisson NMF and ZIFA (not sparse) do not identity the cluster of noisy genes as clearly as sparse pCMF.

**Figure 4:**
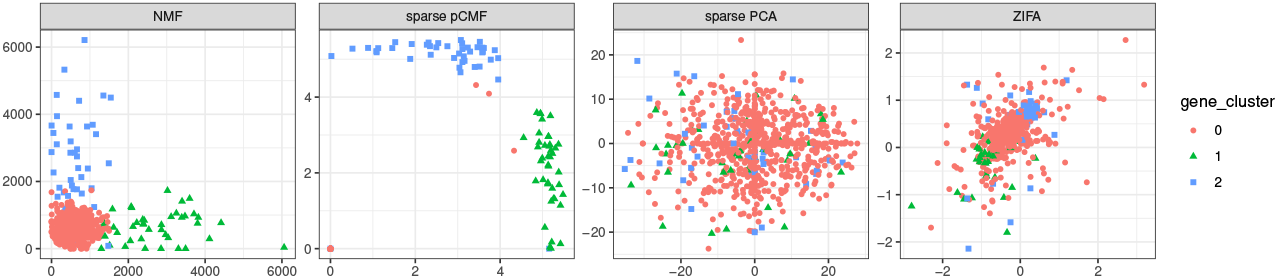
Representation of genes in a subspace of dimension *K* = 2 Here we have 80% of noisy variables, and dropout probabilities around 0.7. The label 0 corresponds to noisy genes.

To quantify the model quality of the different methods on simulated data, we used the deviance associated to each method (c.f. Section 2.1). Figures 1b and 2b shows that the dimension reduction proposed by pCMF has excellent fit to the data regardless the level of dropout or the proportion of noisy genes, as compared with other methods.

Additional comparisons of computational time show that PCA is fastest method (but with low performance), whereas (sparse) pCMF is faster than ZIFA and sparse PCA, with increased performance (cf Supp. Mat. Section S.7).

### 4.2 Analysis of single-cell data

We now illustrate the performance of pCMF on different recent and large single-cell RNA-seq datasets that are publicly availbale: the Baron *et al*. (2016) dataset, the goldstandard and silverstandard datasets used in Freytag *et al*. (2018) (we used the silverstandard dataset 5 which was the largest). We also consider an older and smaller dataset from (Llorens-Bobadilla *et al*., 2015) which is interesting because it discribes a continuum of activation in Neural Stem Cells (NSC). All datasets are available here^3^ with the codes. More details about their origins are given in Supp. Mat. Section S.7.5. We consider large datasets with ~ 1000 or ~ 10000 cells to test the ability of our approach to face the expected increase of data volume in the next couple of years.

We present some quantitative results about clustering and data reconstruction (c.f. Table 1) and the corresponding qualitative results about cell visualization (c.f. Figures 5 and S.7 to S.9 in Supp. Mat.) and gene visualization (c.f. Figures S.10 to S.13 in Supp. Mat.). For each dataset (except the one from Llorens-Bobadilla *et al*., 2015, where we used their pre-filtering), we use the same pipeline, we filter out genes expressed in less than 5% of the cells. In a second step, we remove genes for which the variance heuristic defined in Equation (6) is low. In practice we removed genes for which 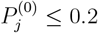. Our idea was to remove uninformative genes, since pre-filtering is crucial (Soneson and Robinson, 2018) in such data, but also to reduce the number of genes to reduce the computation cost, in particular for ZIFA. Then we compare (s)pCMF, PCA, ZIFA and t-SNE. We discarded (s)PCA because the sparse PCA is computationally expensive (c.f. Supp. Mat. Section S.7.3) due to the required cross-validation.

**Figure 5:**
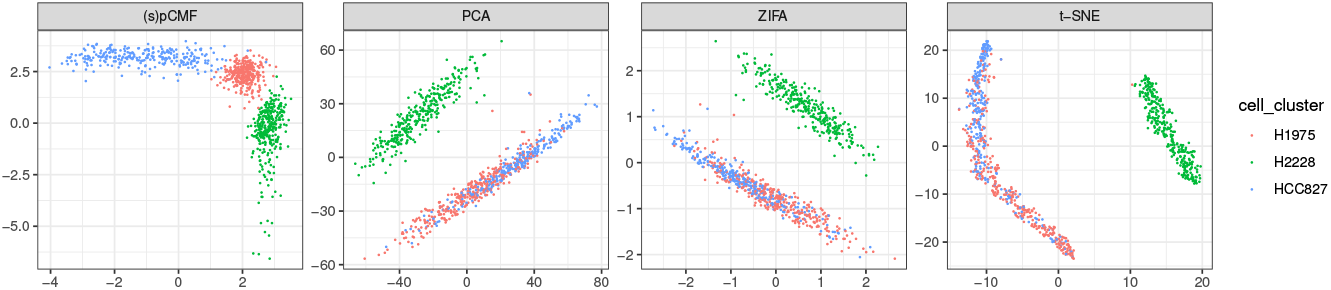
Analysis of the goldstandard scRNA-seq data from Freytag *et al*. (2018), 925 cells, 8580 genes. Visualization of the cells in a latent space of dimension 2.

**Table 1:** Performance of the different methods regarding quality of reconstruction (percentage of explained deviance) and clustering (adjusted Rand Index). Each scRNA-seq dataset is characterized by the number of cells, the number of genes used in the analysis (we specify the original number before the pre-filtering step) and by the number of original groups. The adjusted Rand Index compares clusters found by a *k*-means algorithm (applied to 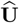 with *k* = nb group) and the original groups of cells.

|  | nb<br>cells | nb genes<br>(before pre-filter.) | prop. 0 | nb<br>group |  | (s)pCMF | PCA | ZIFA | t-SNE |
| --- | --- | --- | --- | --- | --- | --- | --- | --- | --- |
| *Baron et al. (2016)* | 1886 | 6080<br>(14878) | 80.9% | 13 | adj. RI | **21.2%** | 14.3% | 15.4% | 14.2% |
|  |  |  |  |  | %dev | **73.2%** | 41.6% | 53.5% | / |
| *Freytag et al. (2018)*<br>goldstandard | 925 | 8580<br>(58302) | 39.5% | 3 | adj. RI | **81.3%** | 60.1% | 56.8% | 60.5% |
|  |  |  |  |  | %dev | 55.7% | **65.6%** | 48.6% | / |
| *Freytag et al. (2018)*<br>silverstandard 5 | 8352 | 4547<br>(33694) | 86.3% | 11 | adj. RI | 24.2% | 16.2% | 19.8% | **24.8%** |
|  |  |  |  |  | %dev | **70.0%** | 55.1% | / | / |
| *Llorens-Bobadilla et al. (2015)* | 141 | 13826<br>(43309) | 64.8% | 6 | adj. RI | **40.1%** | 25.3% | 38.3% | 29.8% |
|  |  |  |  |  | %dev | **64.4%** | 34.8% | 42.6% | / |

A general empirical property is that clustering accuracy dicreases for all methods when the number of groups of cells increases. However, our approach (s)pCMF produces a better (or as good) view of the cells regarding clustering purpose in every examples, since the adjusted Rand Index is higher (c.f. Table 1). We observe the same trend regarding the quality of the reconstruction since the explained deviance is generally also higher for (s)pCMF. Data visualization is not always clear (c.f. Figures S.7 and S.8), especially when the number of groups is large as in Baron *et al*. (2016) or Freytag *et al*. (2018) silverstandard, however it is possible to clearly identify large clusters of cells in the (e.g. beta cells in Baron *et al*. (2016) or CD14+ Monocyte in Freytag *et al*. (2018) silverstandard) with our method (and some of the others). On the goldstandard dataset from Freytag *et al*. (2018), the difference regarding cells representation between the different approaches is more visual (c.f. Figure 5), where our approach is the only one that is able to distinctly identify the three populations of cells. On the Llorens-Bobadilla *et al*. (2015) dataset, our approach clearly highlights this continuum of activation presented in their paper, which can also be seen with ZIFA, but is not as much clear with PCA and *t*-SNE.

Regarding gene visualization (c.f. Figures S.10 to S.13 in Supp. Mat.), we compare the representation of sparse pCMF to PCA, ZIFA (and not t-SNE since it does not jointly learn **U** and **V**). The interest of sparsity for gene selection is to highlight more precisely the genes that contribute to the latent representation. For each dataset, it is possible to detect which genes are important for each latent dimension: some are null on both (e.g. uninformative genes), some contribute to a single dimension, some contribute to both. This pattern is not as clear with methods that do not implement any sparsity layers.

These different points show the interest of our approach to analyze recent single-cell RNA-seq datasets, even large ones. Empirical properties studied on simulations are confirmed on experimental data: providing a dimension reduction method adapted to single cell data, where the sparsity constraints is powerful to represent complex single cell data. In addition, our heuristic to assume gene importance based on their variance appears to be efficient (*i*) to perform a rough pre-filtering to reduce the dimension and (*ii*) to discriminate between noisy genes and informative ones directly in the sparse pCMF algorithm. In addition, it appears that our method is fast compared to ZIFA for instance, since it takes less than two minutes on the different examples to run (s)pCMF (sparse pCMF + re-estimation), on a 16-core machine, whereas ZIFA can take between 5 and 25 minutes on the same architecture.

## 5 Conclusion

In this work, we provide a new framework for dimension reduction in unsupervised context. In particular, we introduce a model-based matrix factorization method specifically designed to analyze single-cell RNA-seq data. Matrix factorization allows to jointly construct a lower dimensional representation of cells and genes. Our probabilistic Count Matrix Factorization (pCMF) approach accounts for the specificity of these data, being zero-inflated and over-dispersed counts. In other words, we propose a generalized PCA procedure that is suitable for data visualization and clustering. The interest of our zero-inflated sparse Gamma-Poisson factor model is to replace the variance-based formulation of PCA, associated to the Euclidean geometry and the Gaussian distribution, with a metric (based on Bregman divergence) that is adapted to scRNA-seq data characteristics.

Analyzing single-cell expression profiles is a huge challenge to understand the cell diversity in a tissue/an organism and more precisely for characterizing the associated gene activity. We show on simulations and experimental data that our pCMF approach is able to catch the underlying structure in zero-inflated over-dispersed count data. In particular, we show that our method can be used for data visualization in a lower dimensional space or for preliminary dimension reduction before a clustering step. In both cases, pCMF performs as well or out-performs state-of-the-art approaches, especially the PCA (being the gold standard) or more specific methods such as the NMF (count based) or ZIFA (zero-inflation specific). In particular, pCMF data representation is less sensitive to the choice of the latent dimension *K* regarding clustering results, which supports the interest of our approach for data exploration. It appears (through the explained deviance criterion that we introduced) that the reconstruction learned by pCMF better represents the variability in the data (compared to PCA or ZIFA). In addition, pCMF can select genes that explain the latent structure in the data, thanks to a sparsity layer which does not require any parameter tuning.

An interesting direction to improve pCMF would be to integrate covariables or con-founding factors in the Gamma-Poisson model, for instance to account for technical effect in the data or for data normalization. A similar framework based on zero-inflated Negative Binomial distribution was proposed by Risso *et al*. (2017), and it could be extended to our framework of matrix factorization

## Supplementary materials

### S.1 Generalization of explained variance

In the Gaussian framework, we assume that 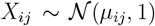 since data are preliminary centered and scaled in PCA. Under the assumptions of independence between observations, the log-likelihood is then in matrix notation:

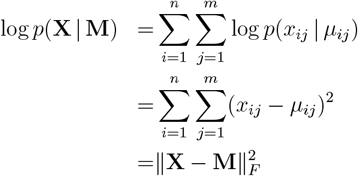

where **M** = [*μ_ij_*] is the matrix of Gaussian expectation and 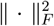 is the squared Frobenius norm. In the generalized PCA framework (Collins *et al*., 2001), we are looking for **U** ∈ ℝ^*n×K*^ and **V** ∈ ℝ^*m×K*^ such that **M** = **UV**^*T*^. Thanks to Eckart and Young (1936) theorem, best **U** and **V** minimizing the Frobenius norm between **X** and **UV**^*T*^ are given by Singular Value Decomposition (SVD) of **X**, and optimal **U** exactly corresponds to the principal components from the PCA, which highlights the link between PCA, SVD and Gaussian framework.

In this Gaussian framework, the explained deviance defined in Equation (3) can be rewritten as

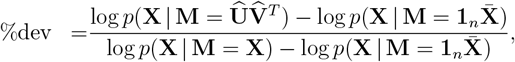

since the saturated model corresponds to **M** = **X** in this case. It follows that

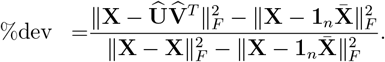

In addition, we have that 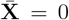 thanks to the pre-centering, and the formulation becomes:

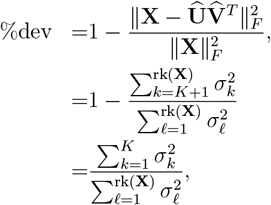

where rk(**X**) is the rank of **X** and *σ*_1_ > … > *σ*_rk_(χ) the singular values of **X** (given by the SVD). The criterion corresponds exactly to the percentage of explained variance computed in PCA. Thus our percentage of explained deviance can be viewed as a generalization of this criterion to other distributions in the exponential family.

### S.2 Identifiability issues

#### S.2.1 Factor order

Gamma-Poisson factor model suffers from an identifiability issue regarding the order of factors. Unlike PCA, the components of model-based factor models are not orthogonal and can not be ordered naturally since the associated likelihood is identifiable up to a permutation of factors. Thus we propose an ordering defined by the cumulative Bregman divergence: 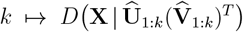. In addition, we mention that the different GaP factor models are not nested when the dimension *K* increases (as in the NMF), thus the factor estimates should be all computed for every choice of dimension *K*, contrary to PCA. sub

### S.3 Scaling effect in GaP factor model

As stated in Dikmen and Févotte (2012), GaP factor models suffer from identifiability issues, due to the scaling of the Gamma prior parameters ***α*** and ***β***. Indeed, considering 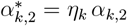 and 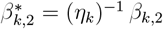 for fixed values *η_k_*, and using the scaling property of the Gamma distribution: if *U_ik_* ~ Gamma(*α*_*k*,1_, *α*_*k*,2_) then 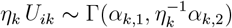.

We show (c.f. below) that the joint log-likelihood regarding **UH**^−1^ and **VH** with **H** = diag(*η_k_*)_*k*=1:*K*_ verifies:

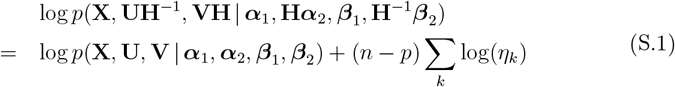

When *n* = *p*, there is an identifiability issue regarding the scaling of the parameters *α*_*k*,2_ and *β*_*k*,2_, because different values lead to the same joint log-likelihood. In such case, a solution will be to fix the scale parameters *α*_*k*,2_ and *β*_*k*,2_ to avoid the scaling effect. When *n* ≠ *p*, the only problem is a potential solution with infinite norm with *α*_*k*,2_ → 0 and *β*_*k*,2_ → ∞ or vice-versa (c.f. Dikmen and Févotte, 2012). When considering zero-inflation or sparsity in the model, Equation (S.1) holds regarding the parameters of the Gamma prior distributions and we have to consider the same precaution. However, in practice we did not encounter such sequence of diverging parameters.

#### Proof of Equation (S.1)

We set, 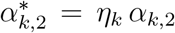 and 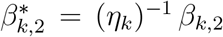 for fixed values *η_k_* > 0. We use the scaling property of the Gamma distribution: if *U_ik_* ~ Gamma(*α*_*k*,1_, *α*_*k*,2_) then *η_k_ U_ik_* ~ Γ(*α*_*k*,1_, (*η_k_*)^−1^*α*_*k*,2_). The joint log-likelihood regarding **UH**^−1^ and **VH** with **H** = diag(*η_k_*)_*k*=1:*K*_ is then:

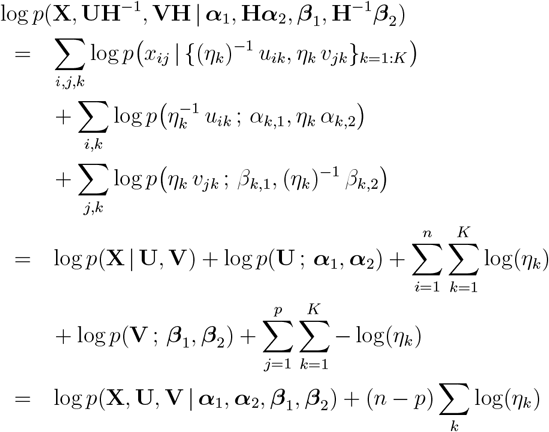

### S.4 Variational inference algorithm

Figure S.1 describes the variational framework (for the GaP factor model) that we extended to develop our approach.

#### S.4.1 Full conditional distributions

In our factor model all full conditionals are tractable. Thanks to the Gamma-Poisson conjugacy, the full conditionals of *U_ik_* and 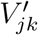 are Gamma distributions. The proof is based on the Bayes rule and the distribution of the latent variables **Z**, that are actually necessary to derive *p*(*U_ik_* | —) and 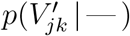. The full conditional of the vector *Z_ij_* is also explicit, being a Multinomial distribution (Zhou *et al*., 2012) when *D_ij_* ≠ 0 and deterministic null when *D_ij_* = 0, i.e. 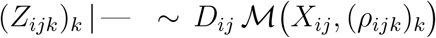. Here the Multinomial probabilities (*ρ_ijk_*)_*k*_ depend on 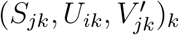, and quantify the prior contribution of factor *k* to the observations *X_ij_*, i.e.

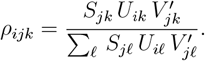

**Figure S.1:**
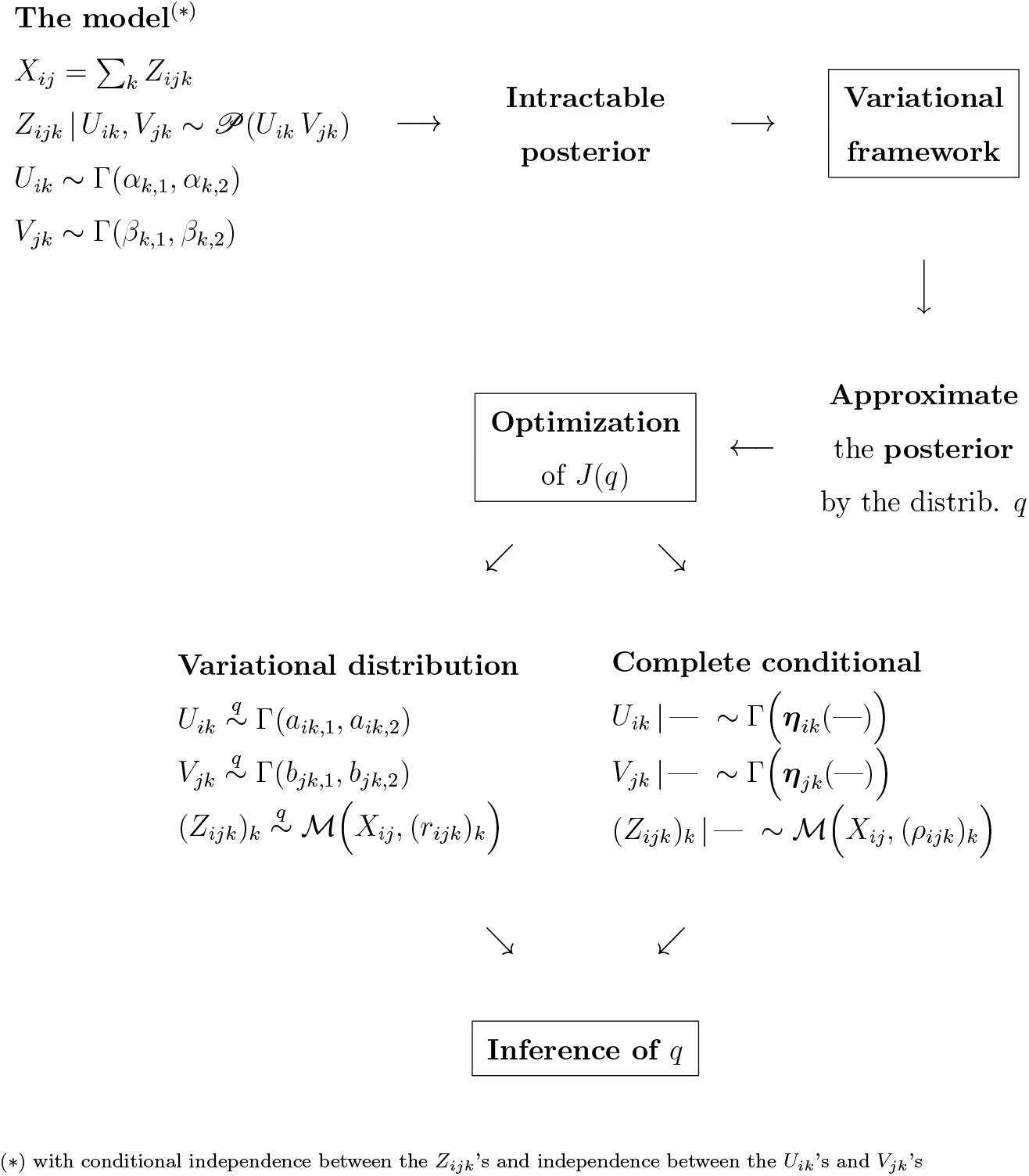
Variational inference to approximate the posterior of the model, based on the optimization of the ELBO that required to derive the full conditional. The notation 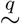 refers to the variational distribution.

This point justifies why the variational distribution is based on the vector **Z**_*ij*_ instead of taking each *Z_ijk_* separately. Note that if the *S_jk_* are null for all *k* or if *D_j_* = 0 (i.e. *X_j_* = 0), the vector (*Z_ijk_*)_*k*_ is deterministic and takes null values. We summarize the full conditionals in the sparse ZI-GaP factor model regarding *U_ik_, V_jk_* and (*Z_ijk_*)_*k*_, that are defined such as:

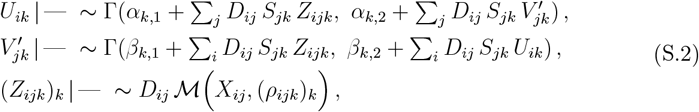

##### Zero Inflation

Regarding the zero-inflation indicators, *D_ij_* is a binary variable, its distribution is either deterministic or Bernoulli. When the entry *X_ij_* is non null, *D_ij_* is certainly equal to one. When *X_ij_* = 0, the full conditional is explicit and the Bernoulli probability only depends on the prior over *D_ij_* and the probability that *X_ij_* is null. It can be formulated as follows:

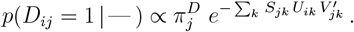

##### Sparsity and variable selection

The sparsity indicator *S_jk_* is also a binary variable and its full conditional is also an explicit Bernoulli distribution. It depends on the prior over *S_jk_* and the probability that gene *j* contributes to the components *k*, quantified by the joint distribution on (*Z_jk_*)_*i*_, thus:

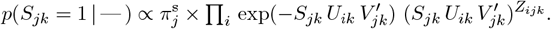

#### S.4.2 Derivation of variational parameters

##### Variational parameters of factors

We derive the stationary point formulation for the variational parameters regarding *U_ik_* and 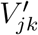, being explicitly (directly derived from the partial derivatives of *J*(*q*)):

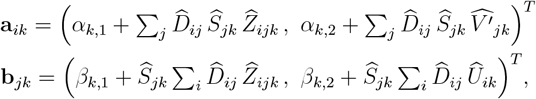

which generalizes formulations from standard GaP factor model (Cemgil, 2009). As for variable *Z_ijk_*, its posterior distribution depends on parameter *r_ijk_* with the relation 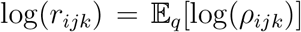. Hence, the variational distribution on (*Z_ijk_*)_*k*_ naturally depends on the selection indicator *S_jk_* (since our model focuses on loadings selection). In particular, the variational parameter *r_ijk_* depends on *S_ijk_*, through a specific term 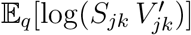 that is computed using the variational distribution of *S_jk_* (a Bernoulli distribution of parameter 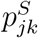). To proceed, we introduce 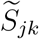, the discretized predictor of *S_ijk_* such that 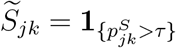, where *τ* is a threshold specified by the user (for instance 0.5). Then, the formulation of the optimal variational parameter *r_ijk_* is approximated by:

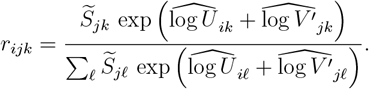

##### Variational dropout proportion

Regarding the zero-inflated probabilities 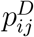, when *X_ij_* ≠ 0, the posterior is explicit since *D_ij_* = 1 with probability one. Hence, only the case *X_ij_* = 0 requires a variational inference. As stated previously, the full conditional is explicit and it is possible to derive and optimize the ELBO (based on the natural parametrization of the Bernoulli distribution in the exponential family). Eventually, 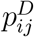 is computed as:

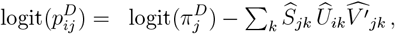

where the Bernoulli prior probability 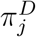 is corrected by 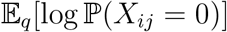 to account for the probability of *X_ij_* being a true zero.

##### Variational Selection probability

Concerning the sparse indicator *S_ijk_*, the natural parametrization of the Bernoulli distribution is based on the logit of the Bernoulli probability. Hence we can write an explicit formulation of the ELBO regarding 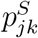 based on the full conditional on *S_jk_*. Following this formulation, the stationary point 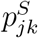 verifies:

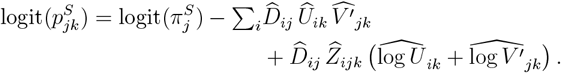

This corresponds to a correction of the Bernoulli prior probability 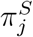, depending on the quantification of the contribution of gene *j* to component *k* in all individuals, i.e. 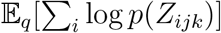.

#### S.4.3 Derivation of prior parameters

The hyper-parameters of priror distribution regarding *U_ik_, V_jk_, D_ij_* and *Sj_k_* are updated within the M-step such that (respectively):

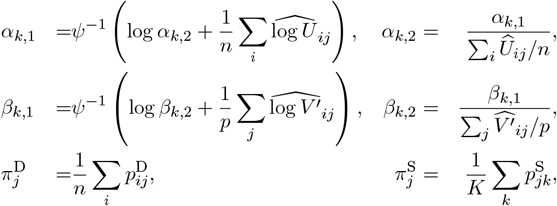

where *ψ* is the digamma function, i.e. the derivative of the log-Gamma function. Its inverse is computed thanks to the method proposed in Minka (2000, appendix C). Recalling that, for a variable 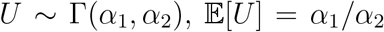 and 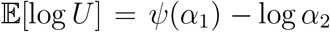, the update rule for the Gamma prior parameters on *U_ik_* corresponds to averaging the moments and log-moments of the variational distribution on *U_ik_* over *i* (similarly for *V_jk_* over *j*). Regarding the Bernoulli prior parameters 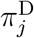, the update rule is also an average of the corresponding variational parameter over *i* (similarly for 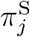 over *k*).

#### S.4.4 Algorithm

Our pCMF algorithm is summarized in Algorithm S.1. In the initialization step, each variational Gamma shape parameter *α*_*ik*,1_ and *b*_*jk*,2_ are sampled from a Gamma distribution (Zhou *et al*., 2012). Each variational Gamma rate parameter *α*_*ik*,2_ and *b*_*jk*,2_ are set to 1 (to avoid scaling effect between shape and rate parameters). Each variational dropout probability 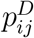 is initialized with the corresponding indicator *δ*_0_(*X_ij_*). Each variational sparsity probability 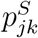 is initialized with the corresponding with the threshold value *τ* ∈ (0,1). In addition, all prior hyper-parameters are initialized following update rules based on variational parameters (defined in Section 3.4 in the paper).

The convergence is assessed by computing the normalized gap between two successive parameter values across iterations. When the updates does not modify the values of the parameters, we can consider that we reach a fixed point and thus the optimum. In addition, to overcome potential issue related to local optimum, the algorithm is run several times with different random initialization and the best seed (regarding the ELBO criterion) is kept.

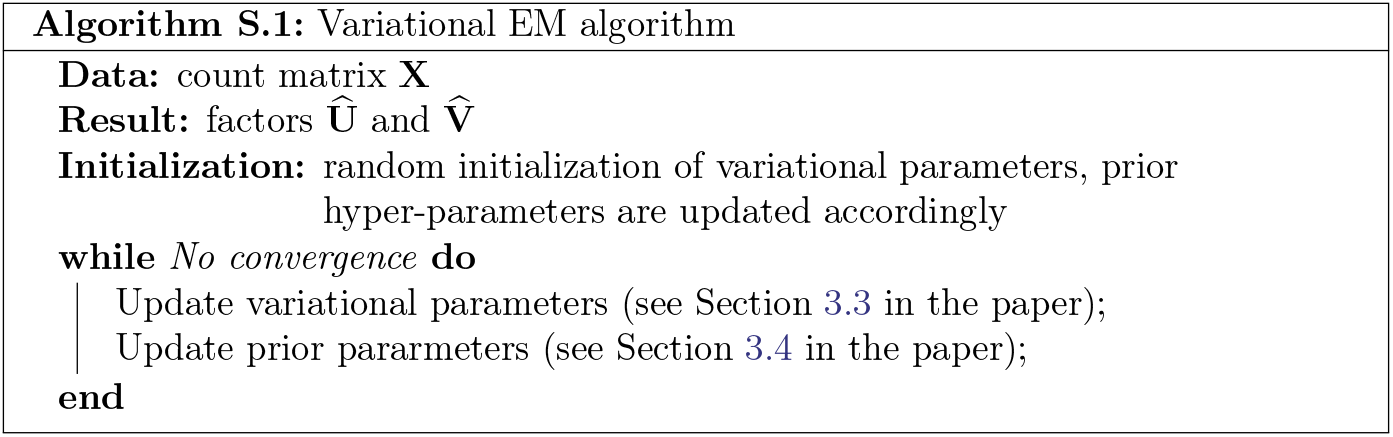

### S.5 Data generation

We set the hyper-parameters (*α*_*k*,1_,*α*_*k*,2_)_*k*_ and (*β*_*k*,1_, *β*_*k*,2_)_*k*_ of the Gamma prior distributions on *U_ik_* and *V_jk_* to generate structure in the data, i.e. groups of individuals and groups of variables.

#### Generation of U

In practice, individuals *i* = 1,…, *n* are partitioned into *N* balanced groups, denoted by 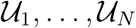. To do so, we generate a matrix U with blocks on the diagonal. Each block, denoted by 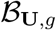 contains *n/N* rows and *K/N* columns. Each entry *U_ik_* in each block 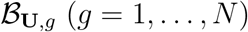 is drawn from a Gamma distribution Γ(1,1/α_g_) with a rate parameter depending on *α_g_* > 0 (different for each group). All entries *U_ik_* that are not in the diagonal blocks of **U** are drawn from a Gamma distribution 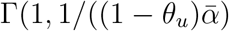 where 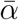 is the average of the *α_g_*’s across *g*, and *θ_u_* ∈ (0,1) quantifies how much the groups of individuals are distinct. Hence, each groups of individuals 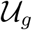 corresponds to a block 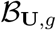. Thus, this generation pattern requires that *K* > *N*. In practice, we fix *α_g_* ∈ {100, 250}, we use θ = 0.5 or 0.8 (for low or high separation respectively) and N = 3 groups of individuals.

#### Generation of V

The question of simulating data based on a sparse representation **V** of the variables in our context of matrix factorization is not straightforward. Indeed, if we impose that some variables *j* do not contribute to any component *k*, i.e. that *V_jk_* is null for any *k*, then Σ_*k*_ *U_ik_ V_jk_* is always null for *i* = 1,…, *n*. Thus, the recorded data entry *X_ij_* will be deterministic and null for any observation *i* (i.e. the *j*^th^ column in **X** will be null). There is no interest to generate full columns of null values in the matrix **X**, since it is unnecessary to use a statistical analysis to determine that a column of zeros will not be informative. This question is not an issue about the formulation of the model, but rather concerns the generation of non informative columns in **X** that will correspond to null rows in the matrix **V**.

To overcome this issue, we use the following generative process. The variables *j* = 1,…,*p* are first partitioned into two groups 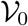 and 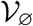 of respective sizes *m*_0_ and *m* − *m*_0_ (with *m*_0_ ≤ *m*). The *m*_0_ variables in 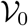 will represent the pertinent variables for the lower dimensional representation, whereas variables in 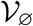 will be considered irrelevant or noise. The matrix **V** will be a concatenation of two matrices **V**^0^ and **V**^∅^:

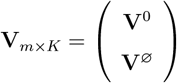

The ratio *m*_0_/*m* sets the expected degree of sparsity in the model. In practice, we generate *m*_0_/*m* from a Beta distribution, so that in average *m*_0_/*m* ∈ {0.2,0.4, 0.6, 0.8} corresponding to different proportions of noisy genes (between 20 and 80% of noisy genes).

To simulate dependency between recorded variables, we generate groups of variables in the set 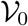 of pertinent variables. We use a similar strategy as the one used to simulate **U**. 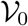 is partitioned into *M* balanced groups, denoted by 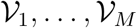. We generate the corresponding matrix **V**^0^ with blocks on the diagonal. Each block, denoted by 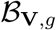 contains *m*_0_/*M* rows and *K/M* columns: Each entry *V_jk_* in each block 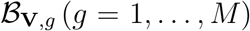 is drawn from a Gamma distribution Γ(1,1/*β*) with a rate parameter depending on *β* > 0. All entries *V_jk_* that are not in the blocks on diagonal are drawn from a Gamma distribution Γ(1,1/((1 – *θ_v_*)*β*), where *θ_v_* ∈ (0,1) quantifies how much the groups of individuals are distinct. Hence, each groups of individuals 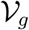 corresponds to a block 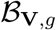. Again, this generation pattern requires that *K* > *M*. In practice, we fix *β* = 80, we use = 0.8 and *M* = 2 groups of variables.

In addition, all *V_jk_* in **V**^∅^ (noisy genes) are drawn from a Gamma distribution Γ(1,1 /((1 – *θ_v_*)*β*), so that 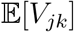 will not be structured according to groups.

#### Generation of X

The data are simulated according to their conditional Poisson distribution in the model i.e. 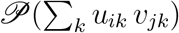. In practice, we want to consider zero-inflation in the model, thus we consider the Dirac-Poisson mixture and simulate *X_ij_* according to the following conditional distribution:

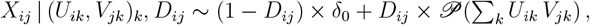

where the dropout indicator *D_ij_* is drawn from a Bernoulli distribution 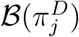, the proportion of dropout events is set by the probability 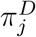. To generate data without dropout events, we just have to set *D_ij_* = 1 for any couple (*i, j*), i.e. 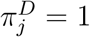 for any *j*.

In practice, we fix *K* = 40, *n* = 100 and *m* = 800 to simulate our data. We generate different level of zero-inflation: 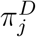 is drawn from a beta distribution so that in average it lies in {0.3, 0.5, 0.7, 0.9}.

### S.6 Softwares

The Poisson-NMF is from the NMF R-package (Gaujoux and Seoighe, 2010), ZIFA from the ZIFA Python-package (Pierson and Yau, 2015), the sparse PCA from the PMA R-package (Witten *et al*., 2009) and t-SNE from the Rtsne R-package (Krijthe, 2015). Computation of adjusted Rand Index was done thanks to the mclust R-package (Fraley and Raftery, 2002).

**Figure S.2:**
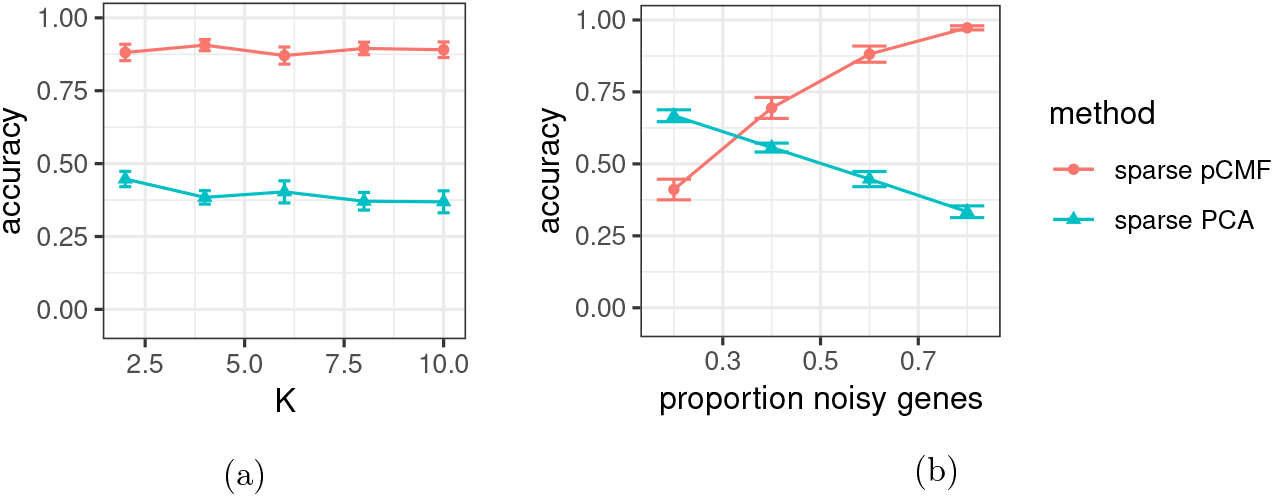
Selection accuracy depending on the dimension *K* (S.2a) with a proportion of noisy genes set to 60% and the proportion of noisy genes (S.2b) with *K* set to 2. Average values and deviation are estimated across 50 repetitions.

### S.7 Additional results

#### S.7.1 Selection accuracy

See Figure S.2.

#### S.7.2 Clustering

See Figures S.3 and S.4. We present results on simulations with different degree of separation between the groups of individuals.

#### S.7.3 Computation time

Figure S.5 shows average computation time for the different methods (pCMF, Poisson-NMF, SPCA, ZIFA) for a single run on a 8-core standard CPU with frequency between 2 and 2.5 GHz. All methods, including ours, have different levels of multi-threading and can benefit from multi-core CPU computations.

Our method sparse pCMF shows comparable computation time as state-of-the-art approaches as Poisson-NMF. The npn-sparse version pCMF is slower but still faster than ZIFA and sparse PCA (because the latter requires a cross-validation step to tune a penalty parameter). t-SNE is slightly faster but requires numerous run with different values for the perplexity parameter (here the timing corresponds to a run for a single perplexity value). The PCA is the gold standard regarding running time thanks to the efficiency of its algorithm based on the Singular Value Decomposition (SVD) algorithm. Packages from where the different methods can be found are detailed in Section S.6.

**Figure S.3:**
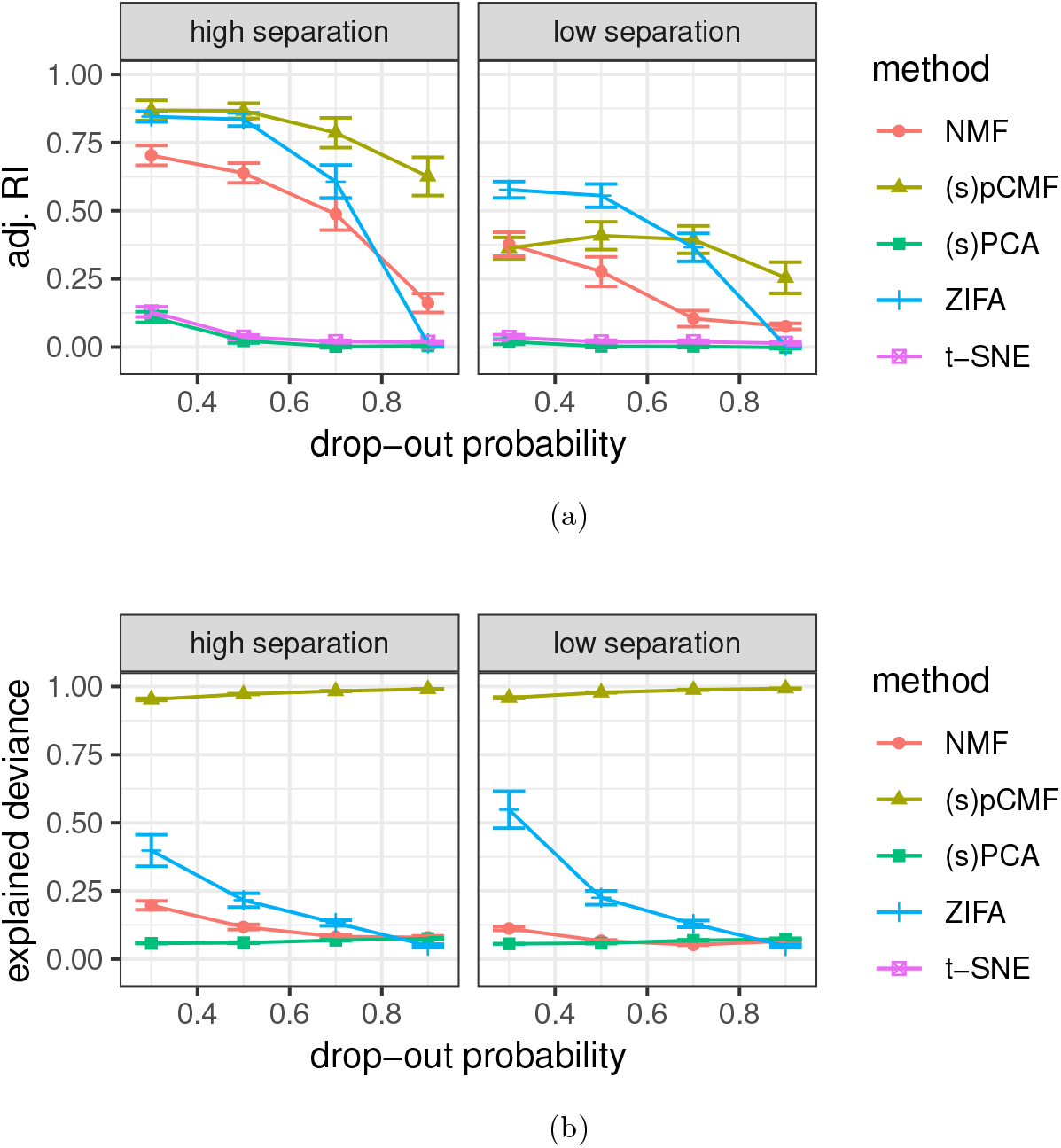
Adjusted Rand Index (S.3a) for the clustering on 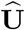 versus the true groups of cells; and explained deviance (S.3b) depending on the probability used to generate dropout events, for different levels of separability between cell groups. Average values and deviation are estimated across 50 repetitions.

**Figure S.4:**
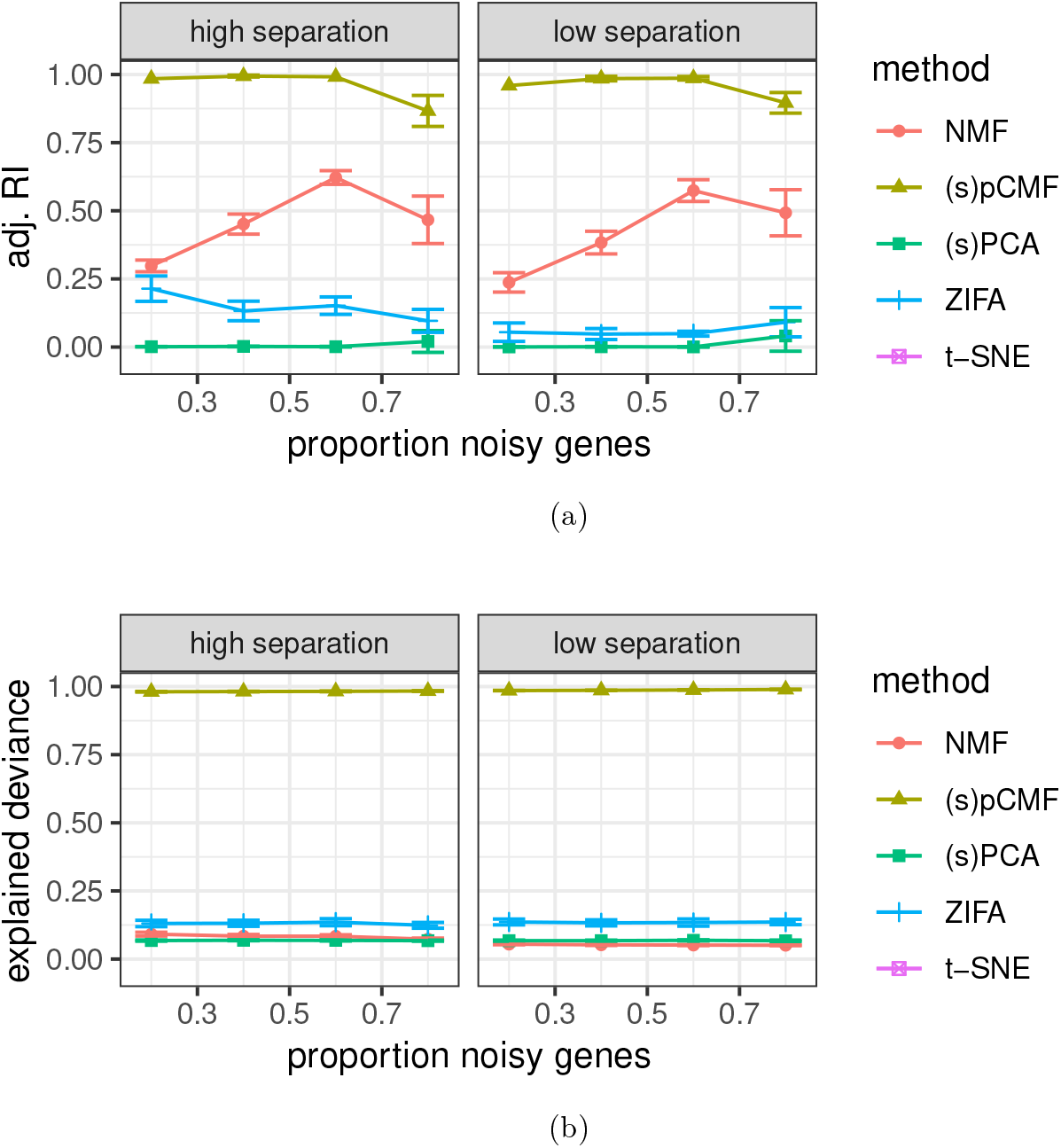
Adjusted Rand Index (S.4a) for the clustering on 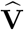 versus the true groups of genes; and explained deviance (S.4b) depending on the proportion of noisy genes, for different levels of separability between cell groups. Average values and deviation are estimated across 50 repetitions.

**Figure S.5:**
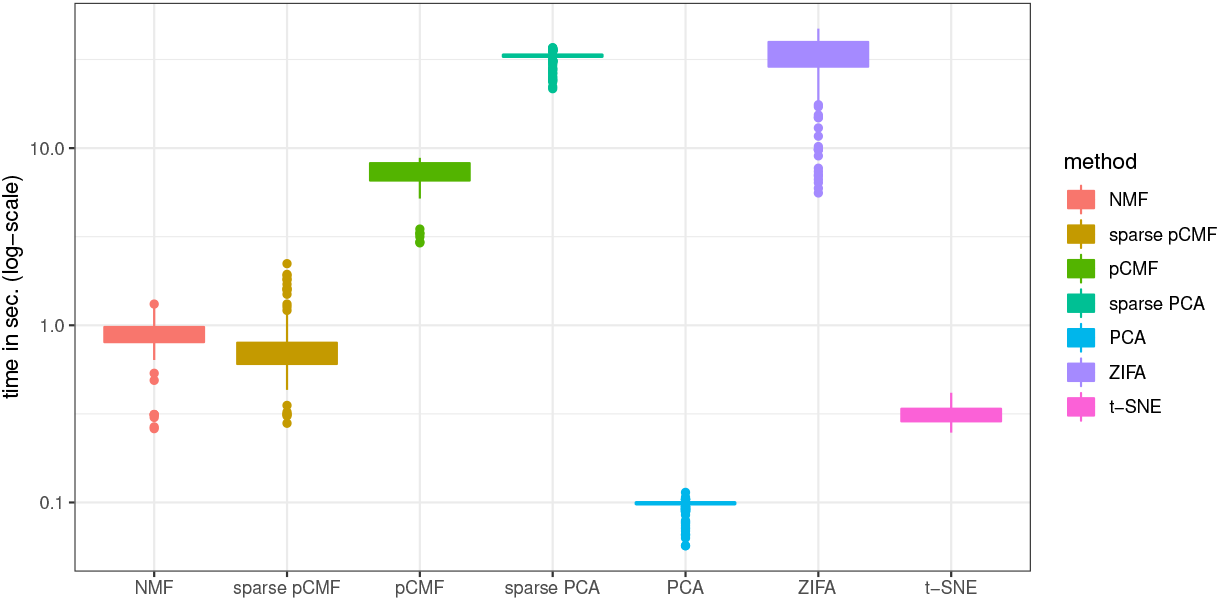
Computation time on 8-CPU core for the different approaches, running time required to analyse simulated data with *n* = 100 individuals and *m* = 800 cells (50 repetitions).

Eventually, we mention that our method is available in an R-package, however our algorithms are implemented in interfaced C++ for computational efficiency.

#### S.7.4 Standard GaP versus our ZI sparse GaP factor model

Figure S.6 illustrates the interest of our zero-inflated sparse Gamma-Poisson factor model compared to the standard Gamma-Poisson factor model, especially in presence of dropout events and noisy genes. Our method pCMF based on our ZI sparse GaP factor model performs as well as the pCMF based on the standard GaP factor model when there is no dropout events in the data, independently from the proportion of noisy genes. In addition, when the level of zero-inflation is higher, we can see that the ZI-specific model outperforms the standard ones, highlighting the interest of our approach.

**Figure S.6:**
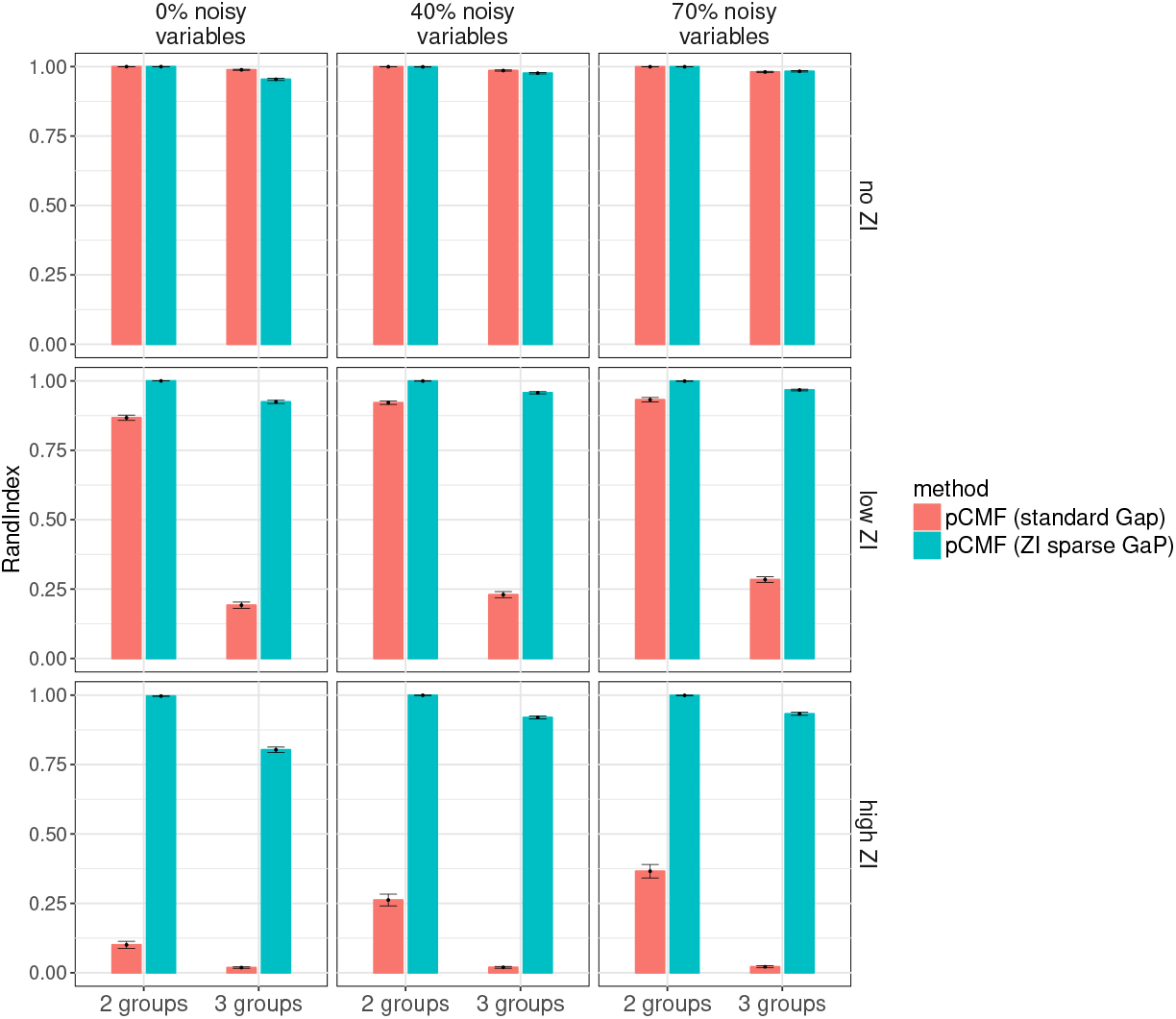
Adjusted Rand Index comparing clusters found by a *κ*-means algorithm (applied to 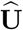 with *κ* = 2) and the original groups of individuals, depending on the number of individual groups in the data, for different levels of zero-inflation and different proportion of noisy variables in the data. The number of components is set to *K* = 10. Data are generated with *n* = 100, *m* = 1000. Average values and deviation are estimated across 100 repetitions.

#### S.7.5 Additional scRNA-seq data analyses

The dataset from Baron *et al*. (2016) is available here^4^. The goldstandard and silverstandard datasets used in Freytag *et al*. (2018) can be found here^5^ (we used the silverstandard dataset 5 which was the largest). The 3 previous datasets are stored based on the `SingleCellExperiment` R package Lun and Risso (2019). The dataset from Llorens-Bobadilla *et al*. (2015) was available as supplementary data of their paper. They kindly shared with us the information about cell tags.

##### S.7.5.1 Llorens-Bobadilla *et al*. (2015)

We illustrate the performance of pCMF on a publicly available scRNA-seq datasets of neuronal stem cells (Llorens-Bobadilla *et al*., 2015). Neural stem cells (NSC) constitute an essential pool of adult cells for brain maintenance and repair. Llorens-Bobadilla *et al*. (2015) proposed a study to unravel the molecular heterogeneities of NSC populations based on scRNA-seq, and particularly focused on quiescent cells (qNSC). In their experiment, qNSC were transplanted in vivo in order to study their neurogenic activity. Following transplantation, 92 qNSC produced neuroblasts and olfactory neurons, whose transcriptome was compared with 21 astrocytes (CTX) and 27 transient amplifying progenitor cells (TAP). The authors used a PCA approach to reveal a continuum of “activation state”, from astrocytes (low activation) to amplifying progenitor cells (TAP).

As stated before, we confront pCMF with other state-of-the-art approaches. The first visual result (c.f. Figure S.9) is that pCMF provides a slightly better representation of the continuum of activation described by Llorens-Bobadilla *et al*. than PCA and t-SNE, which probably reflects a better modeling of the biological variations that exist between activation states. In practice, t-SNE was not able to highlight the different clusters of cells. The results from ZIFA are consistent with pCMF representation, which is a confirmation that the signal of this continuous activation state is strong in these data. These qualitative results are confirmed by clustering quantitative results (c.f. Table 1 in the manuscript). The adjusted Rand Index computed after pCMF and ZIFA are similar and better than PCA.

##### S.7.5.2 Representation of cells

See Figures S.7 to S.9.

**Figure S.7:**
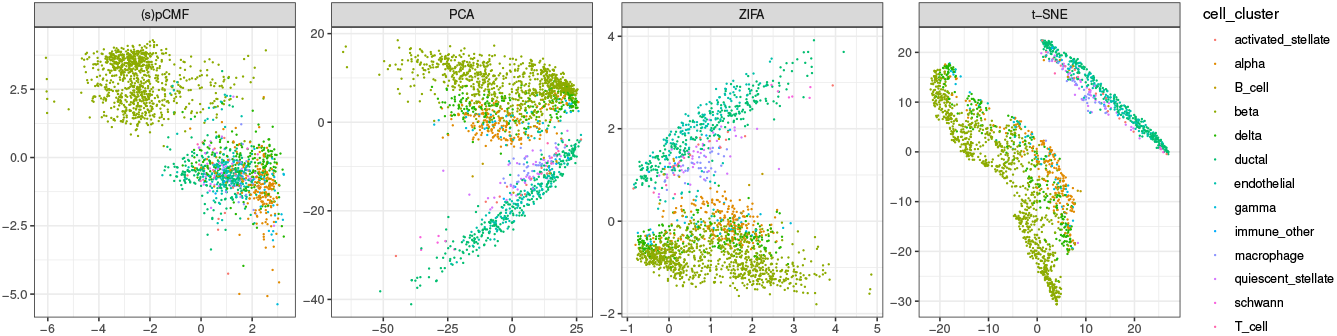
Analysis of the scRNA-seq dataset from Baron *et al*. (2016), 1186 cells, 6080 genes. Visualization of the cells in a latent space of dimension 2.

**Figure S.8:**
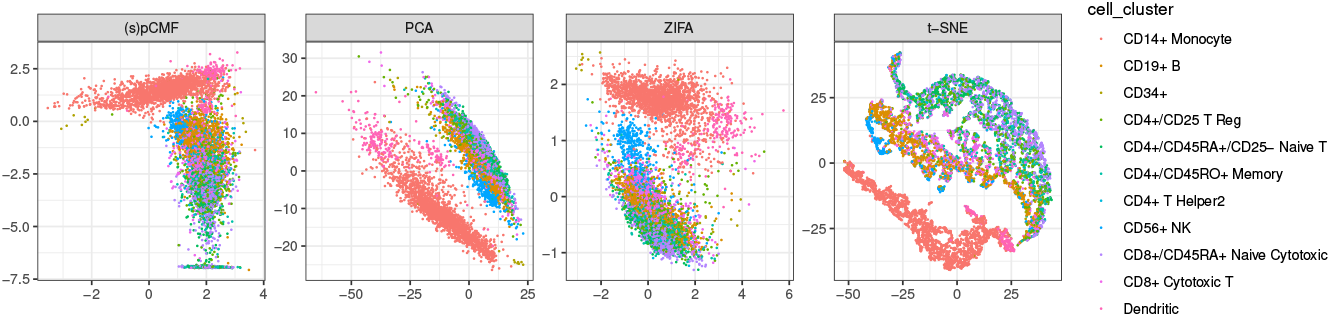
Analysis of the silverstandard 5 scRNA-seq dataset from Freytag *et al*. (2018), 8352 cells, 4547 genes. Visualization of the cells in a latent space of dimension 2.

##### S.7.5.3 Representation of genes

See Figures S.10 to S.13.

**Figure S.9:**
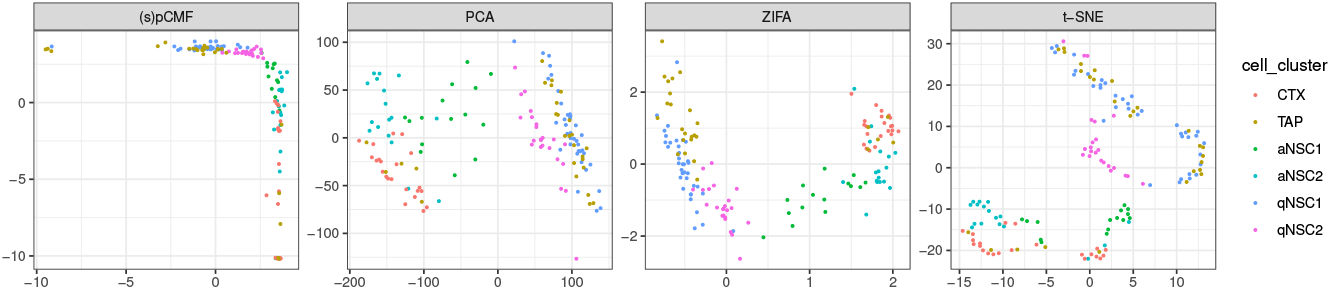
Analysis of the scRNA-seq dataset from Llorens-Bobadilla *et al*. (2015), 141 cells, 13826 genes. Visualization of the genes in a latent space of dimension 2.

**Figure S.10:**
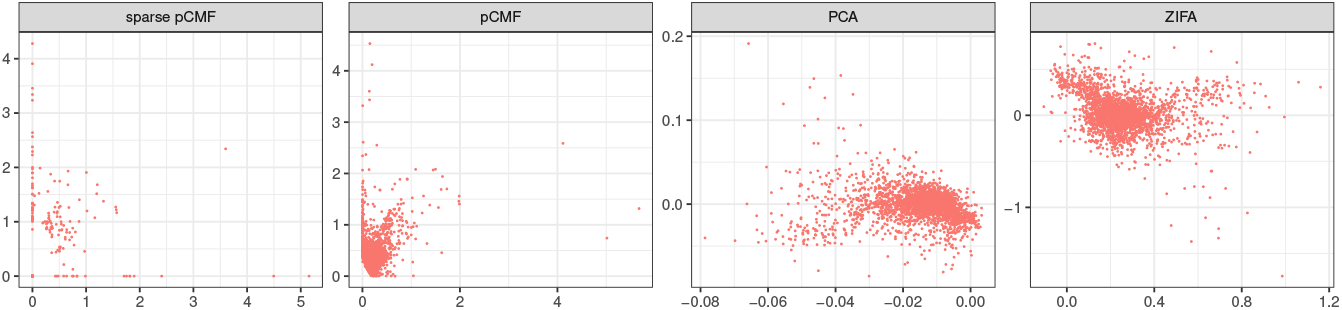
Analysis of the scRNA-seq dataset from Baron *et al*. (2016), 1186 cells, 6080 genes. Visualization of the genes in a latent space of dimension 2.

**Figure S.11:**
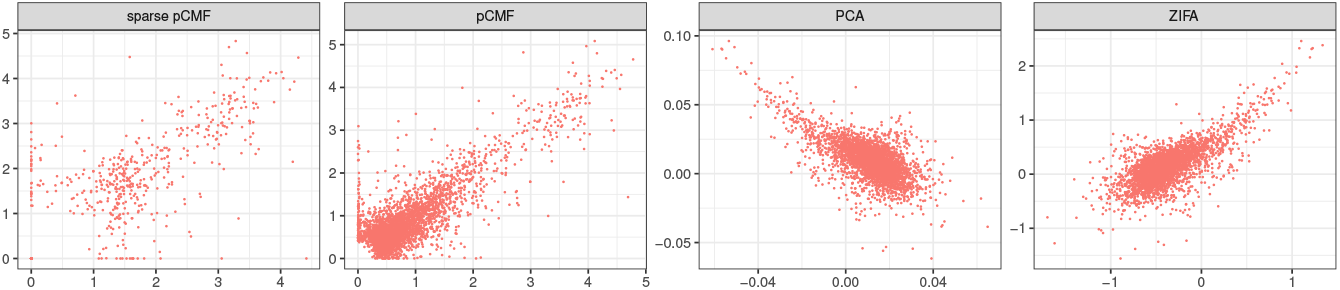
Analysis of the goldstandard scRNA-seq data from Freytag *et al*. (2018), 925 cells, 8580 genes. Visualization of the genes in a latent space of dimension 2.

**Figure S.12:**
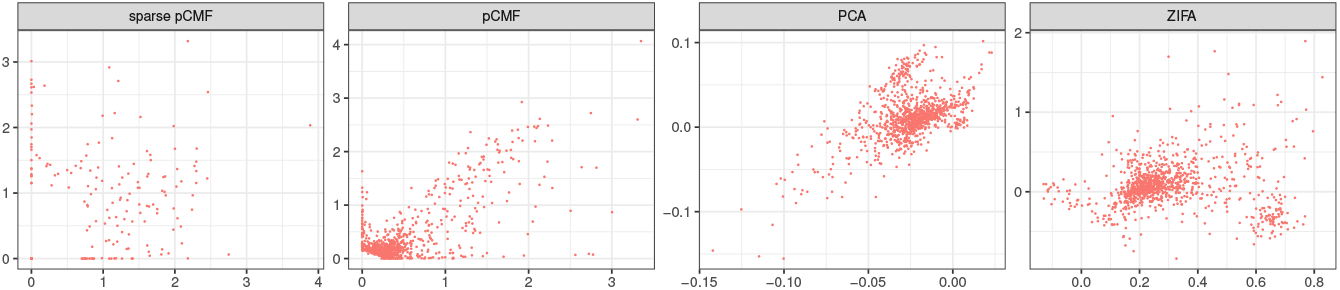
Analysis of the silverstandard 5 scRNA-seq dataset from Freytag *et al*. (2018), 8352 cells, 4547 genes. Visualization of the genes in a latent space of dimension 2.

**Figure S.13:**
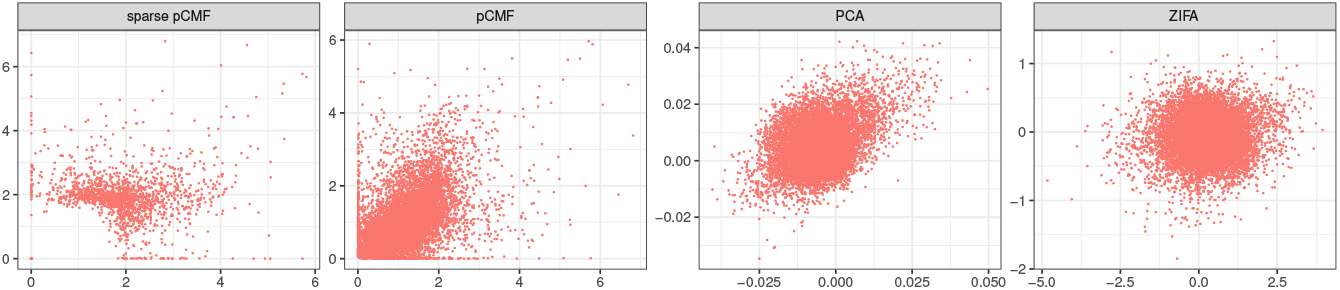
Analysis of the scRNA-seq dataset from Llorens-Bobadilla *et al*. (2015), 141 cells, 13826 genes. Visualization of the genes in a latent space of dimension 2.

## Footnotes

1 https://github.com/gdurif/pCMF

2 https://github.com/gdurif/pCMF_experiments

3 https://github.com/gdurif/pCMF_experiments

4 https://hemberg-lab.github.io/scRNA.seq.datasets/mouse/pancreas/

5 https://github.com/bahlolab/cluster_benchmark_data

